# Dual-Probe Activity-Based Protein Profiling Reveals Site-Specific Differences in Protein Binding of EGFR-Directed Drugs

**DOI:** 10.1101/2023.10.19.562725

**Authors:** Wouter van Bergen, Jan Fiala, Albert J.R. Heck, Marc P. Baggelaar

## Abstract

Comparative, dose-dependent analysis of interactions between small molecule drugs and their targets, as well as off-targets, in complex proteomes is crucial for selecting optimal drug candidates. The affinity of small molecules for targeted proteins is largely dictated by interactions between amino acid side chains and these drugs. Thus, studying drug-protein interactions at an amino acid resolution provides a comprehensive understanding of drug selectivity and efficacy. In this study, we further refined the site-specific activity-based protein profiling strategy, PhosID-ABPP, on a timsTOF HT mass spectrometer. This refinement enables dual dose-dependent competition of inhibitors within a single cellular proteome. Here, a comparative analysis of two activity-based probes (ABPs), developed to selectively target the epidermal growth factor receptor (EGFR), namely PF-06672131 and PF-6422899, facilitated the simultaneous identification of ABP-specific binding sites at a proteome-wide scale within a cellular proteome. Dose-dependent probe-binding preferences for proteinaceous cysteines, even at low nanomolar ABP concentrations, could be revealed. Notably, while both ABPs showed comparable affinities for the EGFR, PF-06672131 had a broader off-target reactivity profile. In contrast, PF-6422899 exhibited higher affinity for the ERBB2 receptor and bound to catalytic cysteines in several other enzymes, which is likely to disrupt their catalytic activity. Notably, PF-06672131 also effectively labeled ADP/ATP translocase proteins at a concentration of just 1 nanomolar. Additionally, analysis of different binding sites within the EGF receptor and the voltage-dependent anion channel 2 revealed secondary binding sites of both probes and provided insights into the binding poses of inhibitors on these proteins. Insights from the PhosID-ABPP analysis of these two ABPs serve as a valuable resource for understanding drug on– and off-target engagement in a dose– and site-specific manner.

## Introduction

Small molecule drugs interact with proteins, affecting protein conformation, activity, and protein-protein interactions [1–4]. Drug-protein affinity is governed by interactions between the drug and amino acid side chains [5]. Even subtle chemical modifications of drugs can markedly alter their affinity toward their protein targets, reshaping the target landscape [6]. Moreover, many of the current drugs target pockets in protein structures, such as ATP-or GDP-binding pockets, that are to some extent conserved over different proteins in the proteome, leading to often undesired off-target binding [7]. Therefore, the investigation of changes induced by subtle differences in the chemistry of the drug on their targets is essential for drug development. Ideally, such investigations should be conducted at an amino acid resolution, allowing for a detailed exploration of the specific interactions between drugs and proteins.

Activity-based protein profiling (ABPP) coupled with peptide-centric enrichment methods enables the site-specific investigation of drug-protein interactions [8–12]. ABPP utilizes activity-based probes (ABPs) to interrogate protein activity and/or (active) site occupancy [13,14]. ABPs consist of a ‘warhead’ to form a covalent bond with target proteins, a recognition element that improves affinity for specific proteins, and a reporter tag to enable visualization or enrichment of targeted proteins [13,15–21]. A peptide-centric enrichment approach enables direct liquid chromatography–mass spectrometry (LC-MS) detection of ABP-bound peptides, resulting in a reduction of detected false positive identifications and a site-specific view of the ABP target landscape [12].

The epidermal growth factor receptor (EGFR), a transmembrane receptor tyrosine kinase and a key player in the regulation of cell growth, is a validated target in cancer therapy [22–25]. Dysregulation of EGFR signaling is linked to several diseases, including cancer, highlighting the importance of effective treatments that target this receptor, ideally with no off-target events [6,22,26,27]. Recently, Lanning et al. introduced two selective Afatinib-derived EGFR-directed ABPs, PF-6422899 (PF899) and PF-06672131 (PF131; a dimethylaminomethyl (DMAM)-modified derivative of PF-6422899), enabling analysis of on– and off-target engagement by ABPP [28]. The comparative protein-centric analysis of the target landscape of these two ABPs showed that DMAM substitution resulted in increased proteome-wide reactivity, which was partly attributed to the prolonged retention times in intact cells [28].

Our study employed a peptide-centric ABPP approach using phosphonate-based enrichment tags (PhosID-ABPP) for the two ABPs in a single cellular proteome [12]. Through their distinctive masses, we obtained a detailed view of the exact localization and relative binding affinities of both ABPs over different concentrations, charting their site-specific binding interactions with proteins.

We were able to quantitatively monitor the binding sites of both probes simultaneously, even at minimal probe concentrations of 1 nM. This dual-probe binding site analysis uncovered diverse binding preferences for PF899 and PF131, even within single proteins. Our dose-dependent evaluation of binding sites allowed us to identify specific and non-specific binding sites for both probes. The charted target landscape provides valuable insight into the effect of small modifications of EGFR inhibitors, enhancing our understanding of EGFR-directed protein-drug interactions.

## Results & Discussion

### Optimization of the mass spectrometry settings on a timsTOF HT for targeted analysis of ABP-modified peptides

This work builds further upon our recent report describing site-specific ABPP using phosphonate handles [12]. To further improve the sensitivity of PhosID-ABPP and facilitate the comparative analysis of multiple ABPs within a complex proteome, we changed to, and optimized the settings on a timsTOF HT mass analyzer for enhanced identification and quantification of ABP-labeled peptides. Four distinct parameters were assessed for the detection of PF131 binding sites in intact A431 cells: 1) the separation and detection range of the trapped ion mobility spectrometry (TIMS) module, 2) the precursor intensity threshold for subsequent collision-induced dissociation (CID) and MS/MS analysis, 3) the collision energy range used for CID, and 4) the inclusion of charge states for MS/MS analysis (Figure 1A).

**Figure 1:**
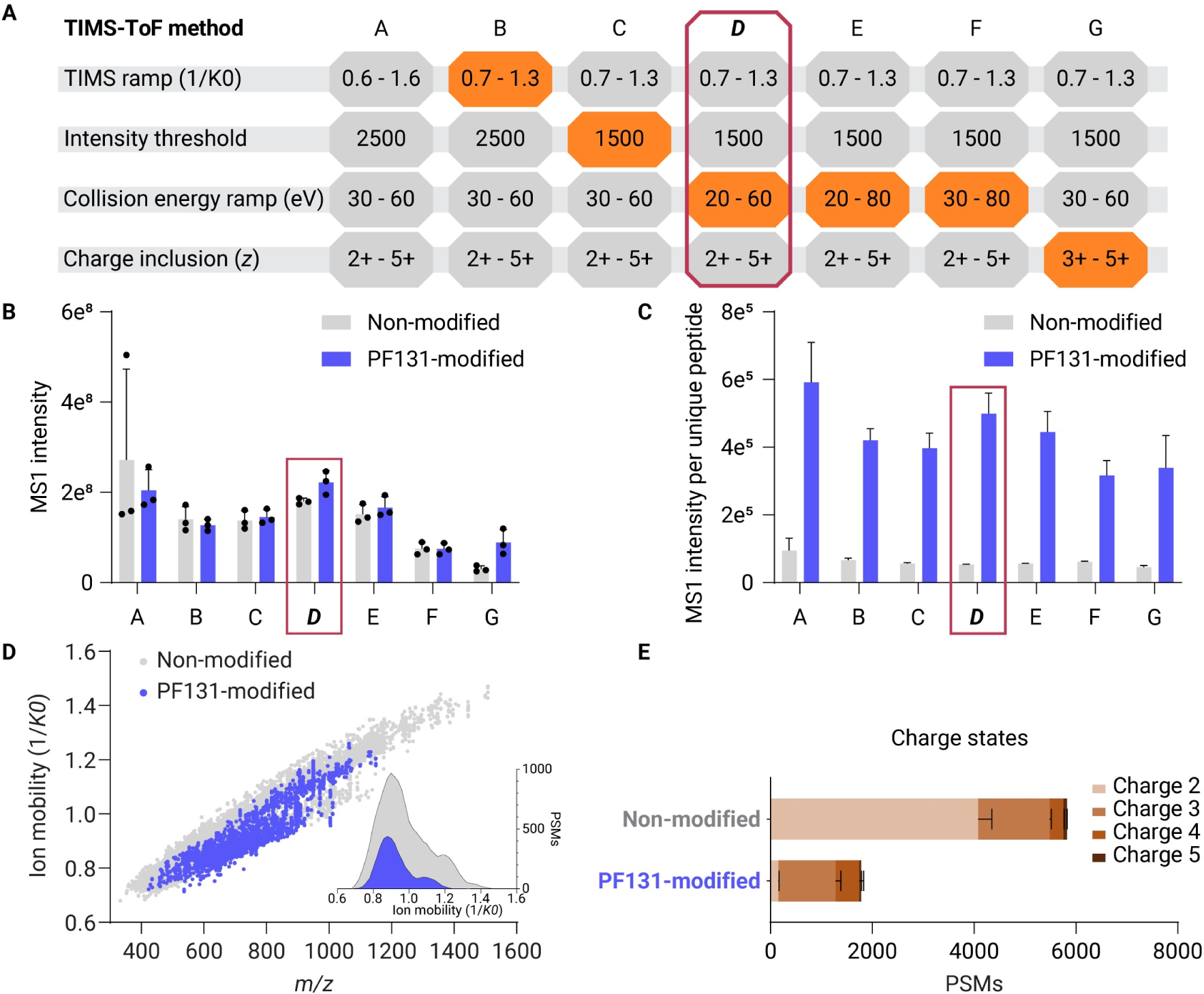
TimsTOF analysis enables efficient detection of activity-based PF131 binding sites. **A)** A schematic of the tested variable parameters in the methods for the detection of PF131-modified peptides. **B)** Summed MS1 intensity of non-modified and PF131-modified peptides using the different methods described in panel A. **C)** Summed MS1 intensity per unique peptide was calculated for both non-modified and PF131-modified peptides. **D)** A plot displaying the ion mobility versus *m/z* of non-modified peptides and PF131-modified peptides detected using method A. The inset displays the distribution of the ion mobility for non-modified (gray) and PF131-modified (blue). **E)** Distribution of peptide-spectrum matches (PSMs) per charge state for both non-modified and PF131-modified peptides.

As a starting point, we used method A, which was optimized for the analysis of non-modified tryptic peptides on the timsTOF HT platform (Figure 1). Analysis using method A resulted in the detection of 1150 PSMs corresponding to 344 unique PF131-labeled peptides, with a summed MS1 intensity of 2e^8^. In addition to these PF131-labeled peptides, 4433 PSMs derived from 2629 unique non-modified peptides were concomitantly identified (summed intensity 2.7e^8^; Figure 1B, Supplementary Figure 1A and B), which on average, indicates a significantly higher intensity for PF131-labeled peptides compared to non-modified peptides (Figure 1C).

Next, we explored the use of ion mobility parameters to enhance the detection of ABP-labeled peptides by isolation/separation of ABP-labeled peptides, adopting a strategy that was previously successful in the analysis of PTMs and cross-linked peptides [29,30]. Comparison of the ion mobility against the *m/z* for PSMs detected by method A revealed a clustering of PF131-modified peptides, spanning an ion mobility range of 0.7 to 1.3 1/*K*_0_. However, the cloud still displayed an overlap with the ion mobility space of non-modified peptides (Figure 1D, Supplementary Figures 1C-G). Notably, the clustering of PF131-labeled peptides can be partially attributed to their higher charge states (Supplementary Figure 1D). The elevated relative abundance of charge states 3+ or higher for PF131-labeled peptides, in comparison to their non-modified counterparts, indicates an additional charge, potentially located on the nitrogen atom of the DMAM group of the ABP (Figure 1E, Supplementary Figures 1C-F). The observed range of the PF131-labeled ion mobility space prompted us to adjust the TIMS ramp from 0.7 to 1.3 1/*K*_0_ for the subsequently explored method (Figure 1A, Methods B-G).

Shortening the TIMS range (method B) resulted in an overall decrease in signal for both PF131-modified and non-modified peptides, possibly indicating a loss of some PF131-labeled peptides at the edges of the TIMS range. Nonetheless, lowering the MS1 precursor intensity threshold (method C) partially mitigated these losses. Further optimization of the ion mobility-dependent collision energy for CID using different ramps (methods C-F) indicated that method D (20 to 60 eV) was the most effective, resulting in the, on average, detection of 1489 PSMs with a summed intensity of 2.2e^8^ from 446 unique PF131-labeled peptides, along with the highest average hyperscore (Figure 1B, Supplementary Figures 1A, B, and H).

Since PF131-modified peptides primarily exist in charge state 3+ or higher, we assessed method G, which exclusively selects precursors with charge states between 3+ and 5+. This approach reduced the detection of non-modified peptides and retained most of the PF131-labeled peptides detected (Figure 1B, Supplementary Figures 1A, B, and C). However, since the PF131-bound peptides derived from the main EGFR binding site could exist in charge state 2 as well, we opted to utilize method D, in the remainder of this work, for the comparative site-specific detection of multiple ABPs in intact cells.

### Comparative dose-dependent profiling of two distinct activity-based probes in a single complex proteome reveals probe-specific characteristics

Employing the optimized method for analyzing ABP-bound peptides, we concurrently investigated the dose-dependent site-specific interactions of two earlier introduced EGFR-directed ABPs, PF131 & PF899, in the context of a complete cellular A431 proteome (Figure 2A).

**Figure 2:**
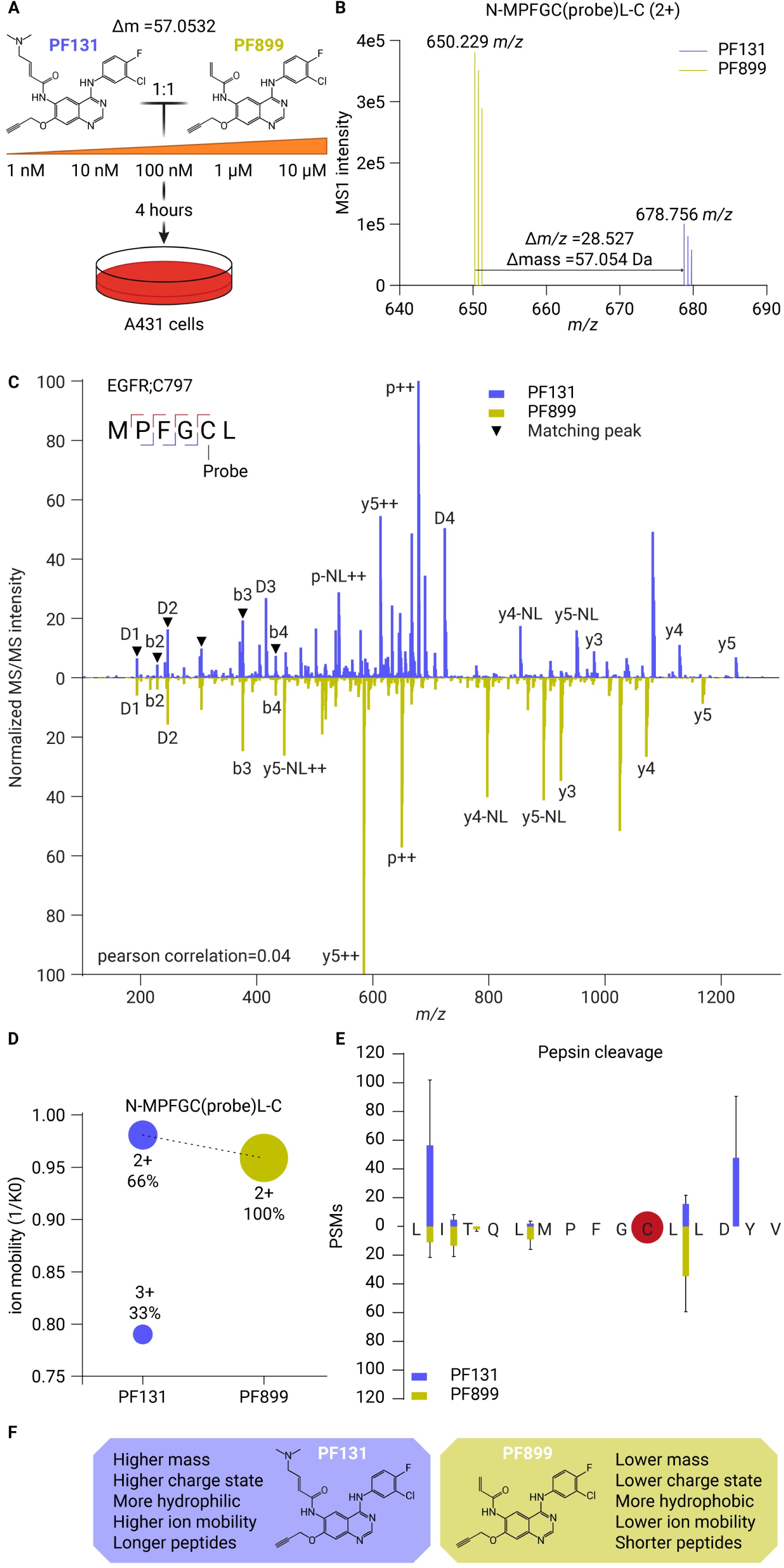
Comparative dose-dependent profiling of two activity-based probes in a single complex cellular proteome reveals probe-specific characteristics. **A)** Experimental design for the comparative dose-dependent profiling of two ABPs simultaneously in intact cells. A431 cells were treated with both PF131 and PF899 (mass difference of 57.0532 Da) mixed 1:1 ([PF131]: [PF899]) in growth medium at different concentrations for 4 hours. PhosID-ABPP analysis was performed to identify the binding sites for both ABPs. **B)** Combined MS1 signal of doubly charged MPFGC(PF131)L (blue) and MPFGC(PF899)L (yellow). The mass difference between the two peptides corresponds to the mass difference between the two ABPs. **C)** Mirror plot of MS/MS spectra of MPFGCL labeled with PF131 (blue) and PF899 (yellow) respectively. Matching and ABP diagnostic ions are marked with a black triangle. b, b-ion; y, y-ion; p, precursor; NL, neutral loss; D, diagnostic ion **D)** Average ion mobility (1/*K_0_*) of MPFGCL bound to PF131 (blue) and PF899 (yellow) in different existing charge states. The size of the dot and the percentage indicate the proportional intensity of the charge state. **E)** Mirror bar graph displaying the peptide-spectrum matches (PSMs) corresponding to the pepsin cleavage site for peptides spanning the EGFR;C797 site bound to PF131 (blue) and PF899 (yellow). The data were obtained from the 10 µM ABP concentration. **F)** Summary of the general different characteristics observed between peptides bound to either one of the two ABPs in the liquid and gas phases.

We selected the non-specific protease pepsin for the proteolysis of ABP-labeled proteomes, based on previous findings demonstrating that pepsin, unlike trypsin, enables the detection of the primary ABP binding site of PF131 on EGFR [12]. Here again, LC-MS analyses identified the covalent attachment of both ABPs to cysteine 797 (EGFR;C797), within the EGFR active site. This attachment was confirmed through the detection of multiple overlapping short peptide sequences encompassing this site, a general advantage when using less-specific pepsin instead of highly specific trypsin. To investigate and illustrate the specific characteristics of both ABPs in the liquid and gas phases, we evaluated the properties of a single peptide backbone (MPFGCL) that was bound by both probes. The mass difference of the peptide bound by the ABPs before fragmentation mirrored the mass difference between the two ABPs (Δ *m/z* = 28.527, Δ mass = 57.054 Da, Figure 2B). Subsequent MS/MS analyses of the MPFGCL peptides bound to the ABPs exposed characteristic differences in fragment ions after CID. MS/MS fragment ions carrying the ABP adduct, or parts thereof consistently showed *m/z* shifts that aligned with the mass difference between the two ABPs. In contrast, fragment ions without the ABP, and diagnostic ions derived from the phosphonate tag matched perfectly between the MS/MS spectra of the differently labeled peptides (Figure 2C).

Analysis of the ion mobility profiles of the two ABPs revealed that PF131 exhibited a minor increase in 1/*K*_0_ for the 2+ charge state of the MPFGCL peptide, which is consistent with its larger molecular size (Figure 2A, D). We observed that MPFGC(PF131)L was detected in both the 2+ and 3+ charge states, comprising 66% and 33% respectively, of the total peak area of the peptide in MS1 scans (Figure 2D). In contrast, MPFGC(PF899)L was detected only as doubly charged ions (Figure 2D). This finding supports our hypothesis that PF131 has an extra positive charge located on its DMAM group, which is absent in PF899. The added hydrophilicity from the DMAM group manifested as a retention time difference between PF131 and PF899, with PF131-modified peptides eluting roughly 11 minutes prior to their PF899-modified counterparts (Supplementary Figure 2C). The differences in retention time, ion mobility, and charge states between the ABP-modified peptides were consistent across all ABP-modified peptides and indicate that they are highly influenced by the properties of the ABP (Figure 2F, Supplementary Figure 2A, B, C).

Interestingly, our analysis elucidated differences in the pepsin cleavage patterns between peptides modified with either one of the two ABPs. Inspection of all EGFR;C797-containing peptides revealed differences in the cleavage pattern at the C-terminus of the peptide sequence (Figure 2E). PF899-modified peptides were predominantly cleaved after the leucine (L798), proximal to the ABP binding site, while the PF131-modified peptides indicated efficient cleavage after the aspartic acid (D800) as well. Importantly, these differences in cleavage specificity affect or even prevent the relative quantification between two specific ABP-bound peptides. Due to these cleavage specificity variations, we suggest that the relative quantification of ABP binding efficacy between probes should be based on peptide populations that cover the specific binding sites. This adjustment is expected to compensate for variations in cleavage specificity. Moreover, given the observed differences in cleavage specificity, we advocate for the use of more and also non-specific proteases in PhosID-ABPP to enhance quantification accuracy by preventing the signal intensity loss from specific ABP-bound peptides incompatible with proteases dependent on a singular specific cleavage site.

Taken together, the timsTOF HT enabled effective detection of ABP binding events from dual-probe-labeled intact cells, revealing significant differences in the liquid and gas phase properties of ABP-labeled peptides, summarized in Figure 2F, which facilitated robust and reproducible detection and quantification of their respective binding sites, further motivating in-depth exploration of these binding sites at lower concentrations.

### Dose-dependent site-specific target landscapes reveal probe-specific target engagement in intact cells

Analysis of the concentration-dependent competitive ABP labeling experiment revealed excellent sensitivity of the site-specific ABPP strategy with PSMs, including those corresponding to PF131-bound EGFR;C797, being detected at a probe concentration as low as 1 nM (Supplemental Data S2-1). To quantify probe-specific concentration-dependent binding events, we generated a library of identified ABP-bound peptides at the 10 µM probe concentration for both PF131 and PF899. Data analysis in Skyline-daily enabled extrapolation of the corresponding MS1 signals at lower concentrations and quantification of ABP-bound peptides across the full concentration range [31]. The MS1 peak areas for all ABP-bound peptides corresponding to specific sites were aggregated and allowed quantification of 613 PF131– and 476 PF899 binding sites at a probe concentration of 10 µM (Figure 3A and B). Substantial off-target binding at a 10 µM ABP concentration was found for both probes. In line with prior research, we observed that PF131 is more reactive than PF899 [28,32]. At lower ABP concentrations, the target landscape for both probes was more restricted. Interestingly, at low ABP concentrations, PF899 appeared to be much more selective with only two targets (EGFR and SOAT1) detected, while 18 distinct target proteins were detected for PF131 at a 1 nM probe concentration. (Figure 3A and B).

**Figure 3:**
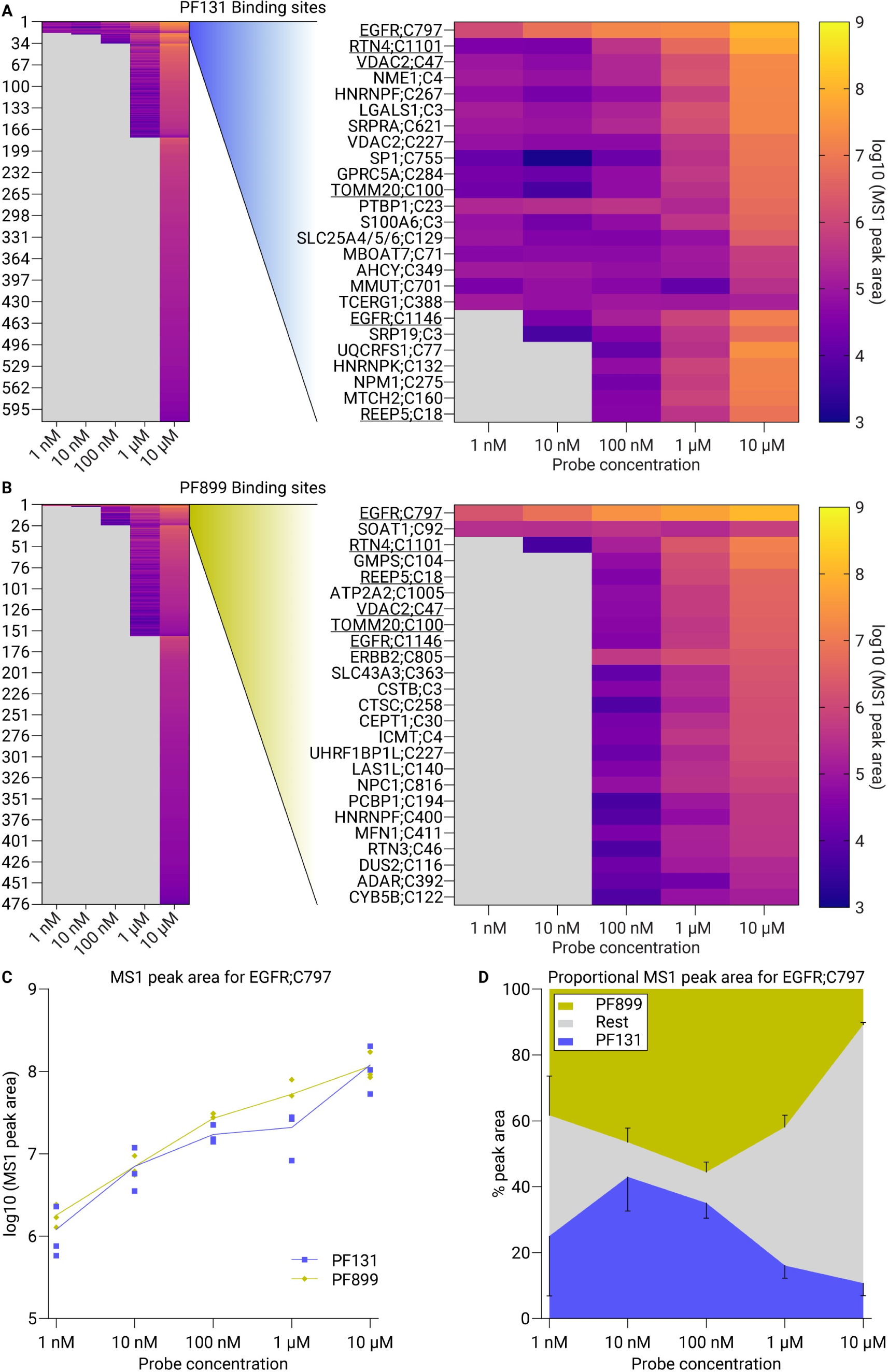
Dose-dependent site-specific target landscape reveals probe-specific target engagement in intact cells. **A/B)** Heatmaps displaying the average aggregated MS1 peak areas of 613 and 476 PF131 and PF899 binding sites at different concentrations in A431 cells, respectively (left segment). The right segments show a zoom of the top-25 binding sites. Underlined sites are top-25 targets for both PF131 and PF899. **C)** Line plot of the aggregated MS1 peak areas for PF131 (blue) and PF899 (yellow) binding to EGFR;C797 across the concentration range. **D)** Density plot of the proportional aggregated MS1 peak areas (*i.e.*, the intensity of the specific site as a percentage of the total probe-labeled intensity in the LC-MS experiment) for the binding of PF131 (blue) and PF899 (yellow) to EGFR;C797 and other sites (gray) across the concentration range.

The concentration-dependent analysis was also conducted in low-EGFR-expressing A549 cells. In this alternative cell line, PF131-binding to EGFR;C797 was also detected and quantified over a broad concentration range. Moreover, the increased promiscuity of PF131 compared to PF899 was recapitulated in this cell line, with 563 and 87 quantified binding sites at 10 µM probe concentration respectively (Supplementary Figure 3A and B). The smaller number of detected binding sites at lower ABP concentrations also applies to A549 cells, with some overlap in the observed binding sites, for instance, PF131-bound VDAC2;C47 and PF899-bound GMPS;C104 at probe concentrations of 10 and 1000 nM, respectively (Supplementary Figure 3C and D). These observations indicate that the ABPP methodology is readily transferable to other cell lines.

As anticipated, EGFR;C797 consistently displayed the highest intensity of all binding sites throughout the concentration range for both ABPs in A431 cells, indicating a high specificity of the probes for their intended primary target (Figure 3A and B). Comparing the concentration-dependent binding of PF131 and PF899 to EGFR;C797 indicated no clear preference for either probe, as both displayed a similar intensity increase across the probe concentration range (Figure 3C). Of note, the proportional intensity of EGFR;C797 (*i.e.* the MS1 peak area of the specified site as a percentage of the total probe-labeled MS1 peak area in the LC-MS experiment) reached its maximum for both probes at 10 and 100 nM (Figure 3D, Supplementary Figure 4A). Off-target labeling increased substantially with concentrations of the probes exceeding 100 nM, mainly caused by PF131’s higher reactivity, suggesting that EGFR-directed inhibitors lacking the DMAM group are superior in selectively inhibiting the EGFR (Figure 3D, Supplementary Figure 4A and B). These results indicate a specificity range of 1 to 100 nM for these probes and suggest binding saturation of EGFR’s ATP pocket above 100 nM, consistent with the approximate IC_50_ of these ABPs (namely, 84 nM for the non-alkynylated derivative of PF899) [28].

In conclusion, the concentration-dependent and ABP-specific analysis of binding site engagement resulted in the detection of over 450 quantified binding sites per ABP. Although both probes show widespread non-specific reactivity in the micromolar range, with PF131 being more promiscuous, they exhibit good specificity for EGFR’s ATP pocket in the nanomolar range. Nevertheless, both probes engage interactions with other binding sites in the nanomolar range, which require further investigation.

### Concentration-response analysis of ABP binding sites elucidates ABP binding preferences in the low nanomolar range

In-depth, concentration-dependent analysis revealed differences in target landscapes between PF131 and PF899 in the nanomolar range. The analysis identified multiple sites with differential binding detection onsets as low as 1 nM, with several sites being unique for individual ABPs, indicating distinct binding preferences.

Our data revealed that ERBB2;C805 is targeted by both PF131 and PF899, with binding commencing at 10 µM and 100 nM respectively (Figure 4A). ERBB2 is a member of the ErbB receptor tyrosine kinase family, to which EGFR also belongs. The binding pocket of its active site cysteine (C805) is homologous to that of EGFR (C797) [33]. Interestingly, while EGFR;C797 was bound by both probes with equal affinity across all concentrations, the MS1 peak areas peptides encompassing ERBB2;C805 revealed that the peak area of PF899 bound to ERBB2;C805 was larger than that of PF131 bound to ERBB2;C805 at a probe concentration of 10 µM (Figure 4B). Furthermore, MS1 peak areas for PF899-bound ERBB2;C805 showed a dose-dependent increase from 100 nM to 10 µM. These results could be extrapolated to the sum of the total peptide population covering ERBB2;C805 for both ABPs, showing an earlier binding onset and significantly higher binding intensity at 10 µM probe concentration of PF899 to ERBB2;C805 compared to PF131 (Figure 4C). Prior research did not indicate a preference for PF899 binding to ERBB2, as both PF131 and PF899 bound with similar intensity at 1 µM [28]. We suspect that this difference may be attributed to variations in study designs. While direct competition between two ABPs at different concentrations in intact cells can reveal binding preferences, separate labeling experiments at a single ABP concentration might not show these competition-induced preferences. The tendency of PF899 to favor ERBB2’s ATP pocket might arise from a decreased affinity when the DMAM moiety is introduced to PF131. This effect might also be seen with other EGFR-directed inhibitors.

**Figure 4:**
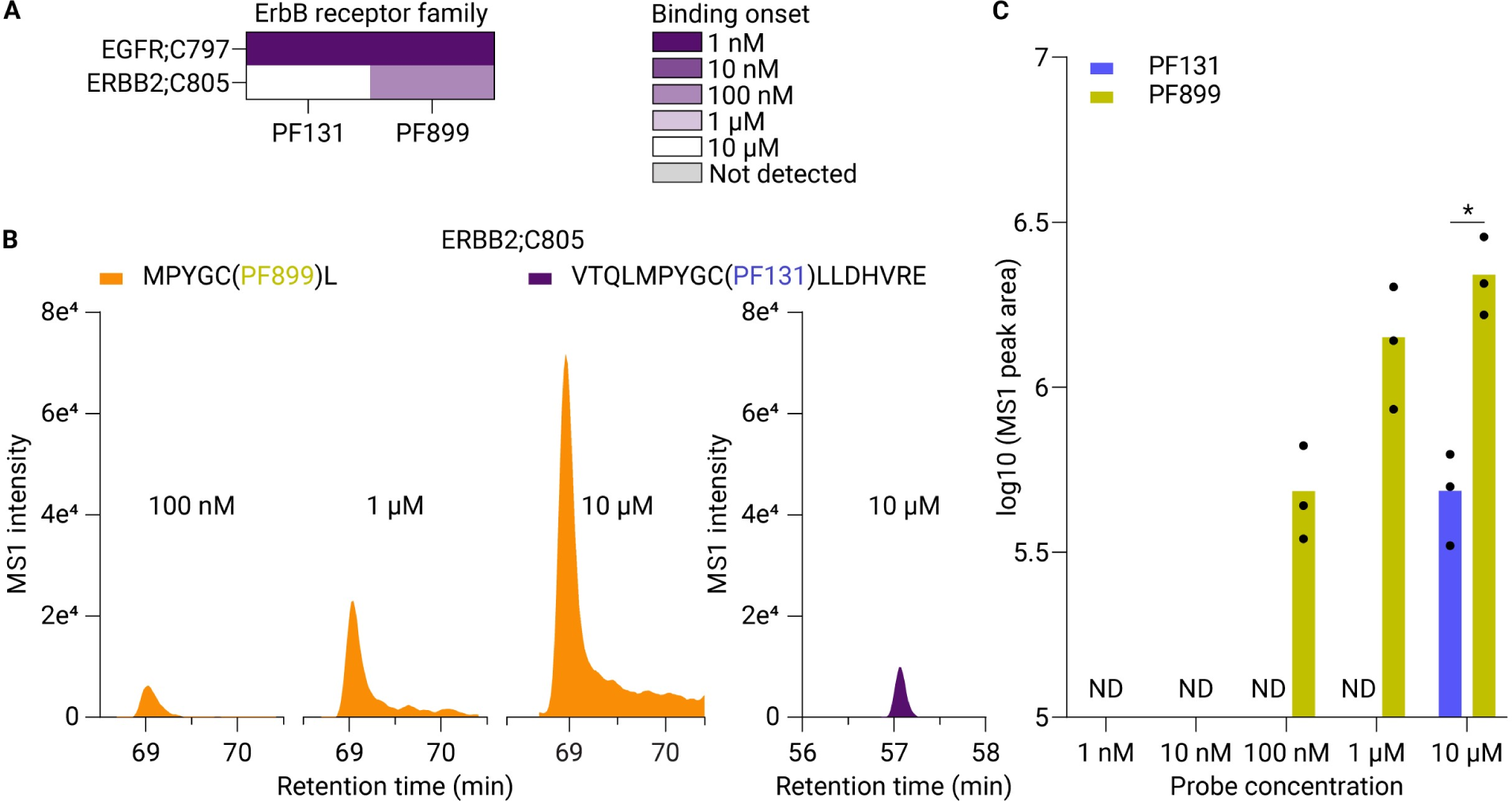
Dose-dependent profiling of ABP binding sites reveals that ERBB2’s active site is relatively favored by PF899. **A)** Binding onset table displaying the ABP concentration at which the ABP initiated binding toward active site cysteines in EGFR and ERBB2. **B)** MS1 peak areas of MPYGC(PF899)L (orange) and VTQLMPYGC(PF131)LLDHVRE (purple) corresponding to both ABPs binding toward ERBB2;C805. **C)** Bar graph of the aggregated MS1 peak areas for PF131 (blue) and PF899 (yellow) binding to ERBB2;C805 across the concentration range. * indicates a significant difference in a paired t-test at the p=0.05 level. ND, not detected.

PF899 binds, in addition to EGFR;C797, to SOAT1;C92 at a 1 nM probe concentration and does not show an increase in labeling intensity with increasing concentrations, suggesting binding saturation at 1 nM. SOAT1’s (Sterol O-acyltransferase 1) cysteine 92 is ligandable by an electrophilic scout fragment, KB05, however the implication of this binding remains elusive [34,35]. ConSurf scores of binding sites show that PF899 targets six conserved cysteines at nanomolar concentrations (The ConSurf score scale ranges from 1 to 9, with a higher score meaning a more evolutionary conserved amino acid) (Figure 5A) [36]. While RTN3;C46 and UHRF1BP1L;C227 are predicted to be structural cysteines and are not known to perform additional functions, three cysteines that are exclusively targeted by PF899 are known as active site cysteines (CTSC;C258, DUS2;C116, and GMPS;C104). Consistent with our data, tRNA-dihydrouridine synthase (DUS2) was previously identified as a preferred PF899 target over PF131, whereas Cathepsin C (CTSC) and GMP synthase (GMPS) were not described as PF899-preferred proteins [28].

**Figure 5:**
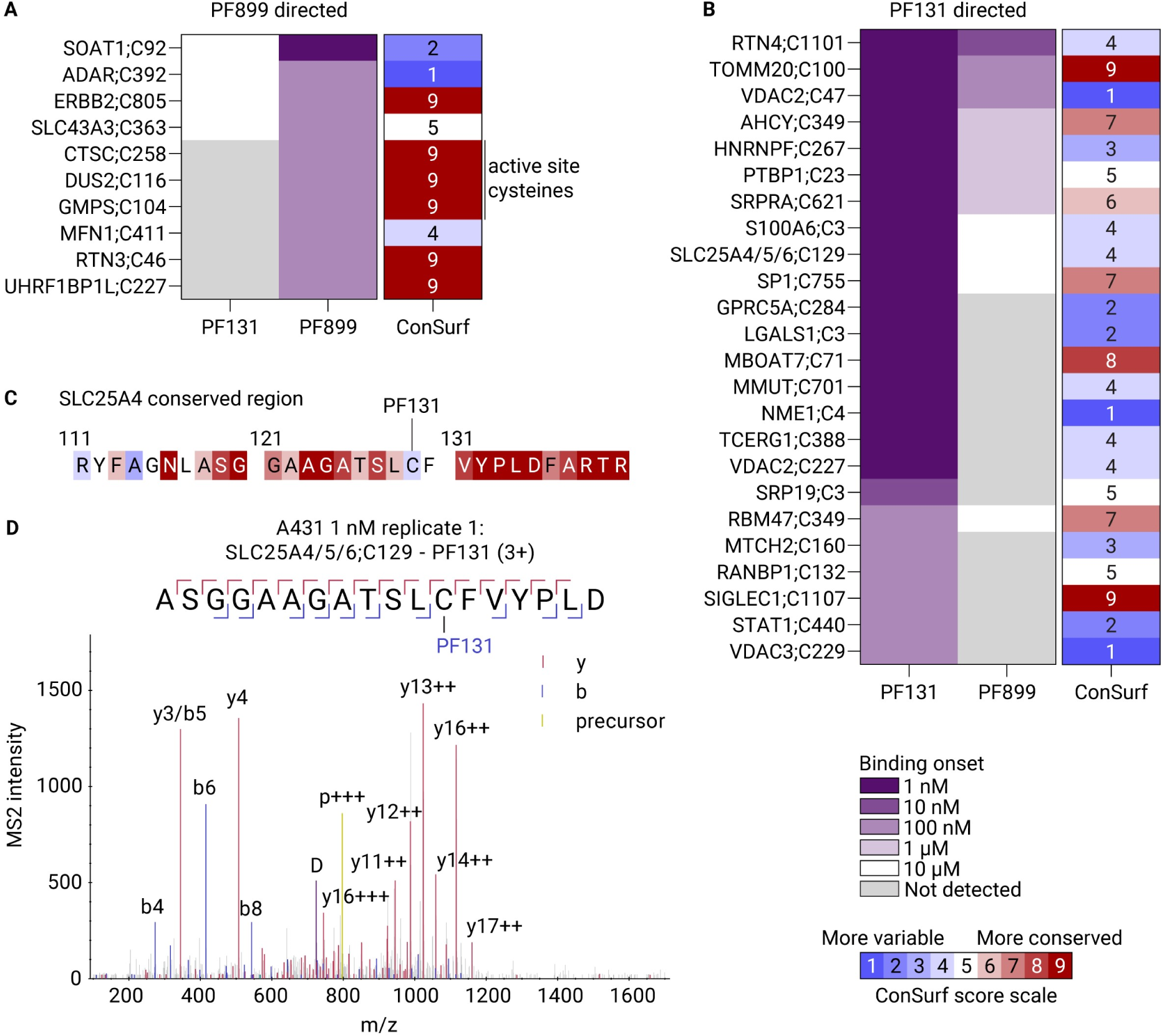
Analysis of off-target ABP binding sites in the nanomolar range elucidates binding preferences for both probes. **A)** Binding onset tables displaying the ABP concentration at which the ABP initiated binding toward the cysteine for PF899-directed (A) and PF131-directed (B) sites. The conservation of the targeted cysteine is expressed in the third column as the ConSurf score. **C)** ConSurf analysis of the region surrounding the PF131-targeted SLC25A4;C129. **D**) Experimental MS/MS evidence for PF131 binding to cysteine 129 on SLC25A4/5/6. b: b-ion, y: y-ion, p: precursor, D: diagnostic ion.

Cathepsin C is a lysosomal exo-cysteine protease, with a critical cysteine in its catalytic dyad, which is targeted by PF899 at a 100 nM probe concentration. This protease plays a key role in inflammatory pathways, activating serine proteases in neutrophil granules through its cleavage function [37]. Cathepsin C is a therapeutic target since its overactivity may lead to disorders caused by hyperreactive neutrophils, such as non-cystic fibrosis bronchiectasis or COVID-19-induced inflammatory diseases [37]. X-ray crystallography of inhibitor-bound CTSC demonstrated covalent binding to cysteine 258, implying that binding at this site renders the protease inactive [38]. Moreover, lysosomal accumulation of small molecule EGFR inhibitors has been proposed to cause increased engagement to cathepsins [6,39]. PF899’s binding to this site and PF131’s lack thereof, may suggest that PF899 accumulates more in the lysosome than PF131, and affect Cathepsin C activity.

GMP synthase, another PF899 target, is a potential target for anticancer and immunosuppressive therapies [40–42]. GMP synthase catalyzes the amination of xanthosine 5′-monophosphate to produce GMP [42]. During catalysis, glutamine is hydrolyzed by cysteine 104, the PF899 binding site, to generate the amino group needed for the amination reaction. The glutamine hydrolysis can be uncoupled from GMP synthesis as other nitrogen sources can also be used. PF899’s binding toward GMPS;C104 shows that PF899 probably inhibits glutamine hydrolysis of GMPS, without interfering with the GMP synthetase function of the enzyme, similar to Acivicin [42]. Our findings suggest that DMAM modification of small molecule inhibitors may affect inhibitor efficacy toward GMPS.

PF131 targets a broader range of proteins at nanomolar concentrations than PF899 (Figure 5B). Several conserved cysteines with ConSurf scores>7 are targeted by PF131 at these concentrations. All of these cysteines are predicted by ConSurf to be buried and perform a structural function in the protein, for example, cysteine 1107 in SIGLEC1 was predicted to form a disulfide bridge with cysteine 1149 [43].

We previously reported PF131’s affinity for Reticulon-4 at cysteine 1101 in intact cells with a concentration of 25 µM [12]. Our current results reveal that RTN4 binding by PF131 can even be detected at a concentration as low as 1 nM, and 10 nM by PF899. MS1 peak areas show that RTN4;C1101 is a preferred binding site for PF131, consistently showing higher intensities than PF899 across the concentration range (Supplementary Figure 5A). The low nanomolar binding of EGFR-directed chlorofluoroacetamides suggests that these inhibitors might cause aberrant ER tubule formation through binding RTN4 and suppress tumor growth at low nanomolar concentrations [12].

Intriguingly, we detected multiple PSMs in all replicates that provided evidence for PF131’s binding toward cysteine 129 on SLC25A4/5/6 at the 1 nM ABP concentration (Figure 5D). These proteins are mitochondrial ATP/ADP transporters, facilitating ADP’s movement into mitochondria and ATP’s export. While PF131 exhibited binding at the lowest concentration, PF899 only binds at 10 µM, suggesting a superior affinity of PF131 for these proteins. Structural insights reveal that cysteine 129 is positioned within an alpha helix spanning the mitochondrial membrane, which, in combination with other helices, forms a pore for ATP in the M-state and ADP in the C-state [44]. Though cysteine 129 is not a conserved residue, with a ConSurf score of 4, the surrounding region is highly conserved and involved in binding ATP or ADP (Figure 5C). Both M– and C-states can be inhibited by bongkrekic acid and carboxyatractyloside respectively, and can cause severe toxic effects at low concentrations [45,46]. PF131’s binding to mitochondrial ATP/ADP transporters might disrupt their function, leading to adverse events. The DMAM moiety in EGFR-directed inhibitors appears to provide improved affinity toward these antiporter proteins and this valuable information could be used when designing novel drugs for ATP/ADP antiporters.

Collectively, our findings indicate pronounced disparities in the binding affinities of PF131 compared to PF899 at nanomolar concentrations, providing insights into the effects of DMAM modification of small molecule drugs might have on their drug-protein interactions.

### Analysis of multiple probe binding sites within individual proteins

Site-specific analysis facilitates the discovery of multiple ABP binding sites within individual proteins. Within EGFR, five distinct ABP binding sites were identified in addition to the anticipated target, C797 in the ATP binding pocket. Four of these sites are localized in the cytosolic domain (Figure 6A). The primary binding site (C797) and cysteine 775, which are both solvent-accessible cysteines localized in the ATP pocket, display substantial disparities in their measured labeling intensities [47]. ABP-modified cysteine 797 can be detected at concentrations as low as 1 nM by both probes and consistently displays higher labeling intensities compared to C775 throughout the whole concentration range. This observation is in line with the binding pose of covalent chlorofluoroacetamide-based EGFR inhibitors, with the Michael acceptor situated near cysteine 797 [33,47]. Except for cysteine 1146, binding to EGFR cysteines other than C797 is only detectable at ABP concentrations of 1 µM and above. C1146-binding by PF131 and PF899 was already detected at 10 and 100 nM respectively. This pattern suggests differential binding affinities across different EGFR cysteines, with both ABPs demonstrating a relatively efficient binding to cysteine 1146 compared to the other non-primary binding sites in EGFR. Relatively efficient binding to cysteine 1146 may be potentially explained by increased solvent accessibility and/or the interactions that the ABP establishes with the protein near this site.

**Figure 6:**
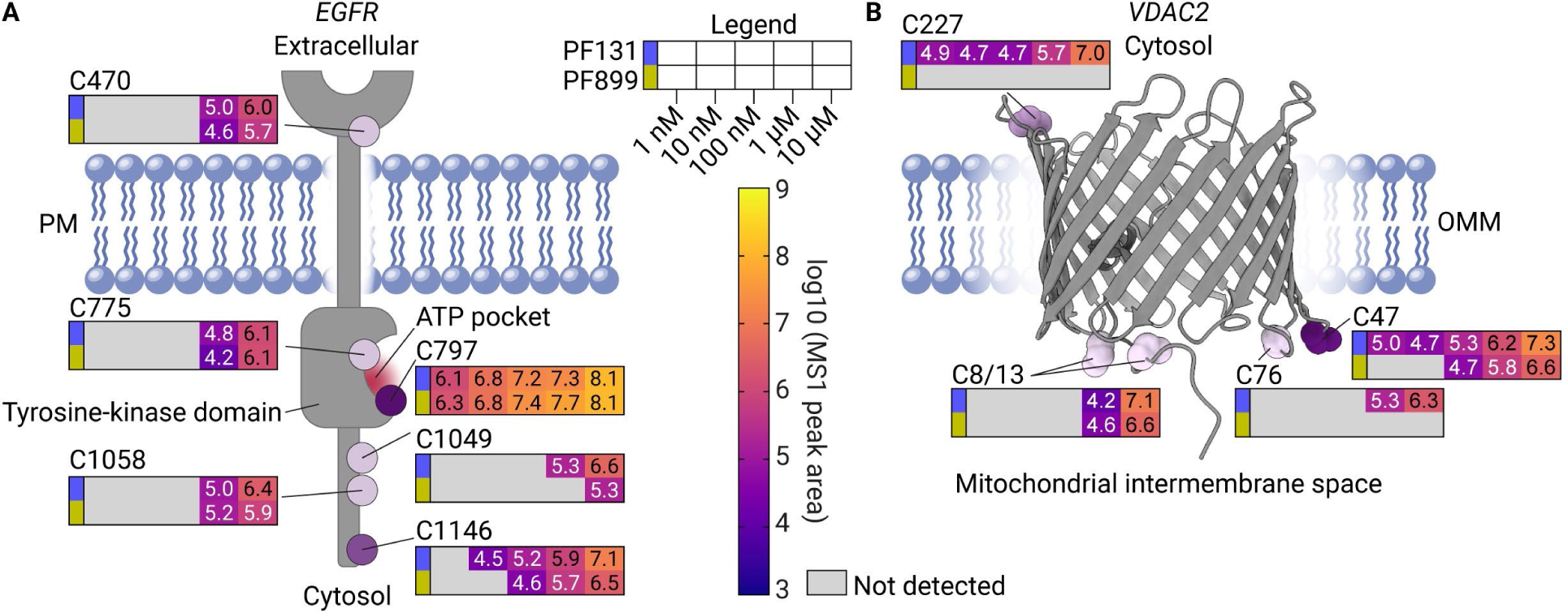
Binding preferences of the two probes within individual proteins. **A)** Topology of EGFR displaying the average aggregated log10 MS1 peak areas at different ABP concentrations on each binding site for PF131 (blue) and PF899 (yellow). PM; Plasma membrane. **B)** Structural model for porcine VDAC2 displaying the average aggregated log10 MS1 peak areas at different ABP concentrations on each binding site for PF131 (blue) and PF899 (yellow), adapted from Leung *et al.*, 2021, PDB: 7NIE [49]. OMM, Outer mitochondrial membrane.

Additionally, voltage-dependent anion channels 1, 2, and 3 were found to exhibit ABP binding sites, with voltage-dependent anion channel 2 (VDAC2) exhibiting five distinct probe binding sites (Figure 6B, Supplementary Figure 6). In line with previous studies on cysteines in VDAC isoforms, totally reduced and redox-active cysteines in VDAC,1 2, and 3 are primarily targeted by both probes [48]. Mapping the bound cysteines on an electron microscopy model of porcine VDAC2 shows that most of these cysteines are located in the mitochondrial intermembrane space (Figure 6B) [49]. Notably, cysteine 47 displayed the highest labeling intensity by both ABPs, with PF131 and PF899 binding being detected at 1 and 100 nM ABP concentrations, respectively (Figure 6B, Supplementary Figure 5B). This suggests an ABP binding pose with the Michael acceptor of the ABPs proximal to C47. Furthermore, CysDB was used to assess the reactivity of cysteines in VDAC2. This analysis revealed that VDAC2 has two known hyperreactive cysteines, namely C210 and C227 [34,35,50]. Only C227 bound to PF131 was detected in our experiments, which points toward labeling of the cysteines driven by ABP affinity for specific VDAC2 sites, rather than just cysteine reactivity.

In summary, these findings demonstrate the binding of both probes at different cysteines within individual proteins, revealing secondary binding sites and providing insights into preferred binding orientations of ABPs within proteins. Importantly, differentiating between primary and secondary binding sites of small molecule drugs in proteins is unattainable without site-specific ABPP. Furthermore, binding to secondary sites within proteins can elicit functional consequences, making it an essential aspect to consider during drug binding evaluation by ABPP.

## Conclusions

In this study, we extended upon our earlier research on site-specific activity-based protein profiling using phosphonate handles, integrating simultaneous dose-dependent competition of two activity-based probes in intact cells. To identify optimal settings for the detection of ABP-bound peptides on the timsTOF HT, we leveraged unique characteristics of ABP-bound peptides compared to unmodified counterparts to enhance their detection. Furthermore, we discovered specific properties for each probe across multiple dimensions within a single LC-MS/MS run, including reversed-phase liquid chromatography, ion mobility separation, enzymatic proteolysis, and collision-induced dissociation. The detailed deconvolution of ABP binding sites allowed for the relative quantification of binding intensities to cysteine residues across the total proteome for two EGFR-directed probes. While both probes showed similar affinities for the EGF receptor, PF-06672131 displayed a broader off-target profile in all concentrations. PF-6422899 displayed a higher affinity for the ERBB2 receptor and bound specifically to catalytic cysteines in CTSC, DUS2, and GMPS, likely disrupting their enzymatic activity. Our analysis revealed that PF-06672131 might affect mitochondrial ADP/ATP translocase activity from a concentration of just 1 nanomolar by binding to SLC25A4/5/6. Lastly, unlike a protein-centric enrichment approach, the analysis of different binding sites of both probes within single proteins, here demonstrated on EGFR and VDAC2, may aid in the identification of secondary binding sites or predict the binding poses of inhibitors. Insights from the PhosID-ABPP analysis of these two ABPs serve as a valuable resource for understanding drug on– and off– target engagement in a dose– and site-specific manner, elucidating the effect of DMAM addition on EGFR-directed inhibitors and contributing to the advancement of drug development efforts.

## Experimental Procedures

### Cell Culture

A431 and A549 cells (CRL-1555 and CCL-185, ATCC) with a passage number below 20 were cultured in growth medium [(Dulbecco’s modified eagle medium (Gibco) supplemented with 10% fetal bovine serum (HyClone GE) and 100 units/ml penicillin-streptomycin (Gibco)]. Cells were grown in a humidified atmosphere with 5% CO_2_ at 37 °C in T175 flasks (Greiner). Cells were split twice a week by washing with Dulbecco’s phosphate buffered saline (DPBS, Lonza) and treated with 0.05% Trypsin-EDTA (Gibco) for cell detachment. After detachment, trypsin was quenched by adding growth medium. 1/10 of the cell suspension was taken and grown with fresh growth medium in a new T175 flask.

### Activity-based probe incubation in cell culture

5e^6^ cells were plated in 15 cm plates (Greiner) 1 day before probe incubation and kept in a humidified atmosphere with 5% CO_2_ at 37 °C. The growth medium was replaced by treatment medium [growth medium with corresponding concentration of PF-06672131 and/or PF-642899 (Sigma-Aldrich)] and incubated at 37 °C, 5% CO_2_ for 4 h. Cells were washed with ice-cold DPBS, harvested using a cell scraper in 1 mL ice-cold DPBS. Then the cell suspension was spun down at 400 g for 5 min, and the supernatant was aspirated. The cell pellet was snap-frozen in liquid nitrogen and stored at −80 °C for later use.

### Cell Lysis

Cell pellets were lysed in 500 μl Lysis buffer per 15 cm plate, consisting of 50 mM HEPES (Sigma-Aldrich, pH 7.5), 0.5% NP-40 (Applichem), 0.2% SDS (Gen-Apex), 2 mM MgCl_2_ (Sigma-Aldrich), 10 mM NaCl (Merck), 1X protease inhibitor cocktail (Roche), and 0.5 µL/mL Benzonase (Millipore). Cell lysates were incubated at room temperature for 15 minutes to allow DNA cleavage. Cell debris and DNA were spun down for 30 min at 20,567g at 16 °C. The supernatant was collected, and the protein concentration was determined by a bicinchoninic acid assay (Thermo Fisher Scientific).

### Bioorthogonal Chemistry Reactions for Proteomics

The copper(I)-catalyzed azide-alkyne cycloaddition (CuAAC) was performed on 4 and 2.5 mg protein lysates for timsTOF optimization and dual-probe analyses respectively in 2 M urea (Merck) in 1× 50 mM HEPES (pH 7.5). CuAAC components were added in the following order: 5 mM tris(3-hydroxypropyltriazolylmethyl)amine (Lumiprobe), 2.5 mM CuSO_4_ 5·H_2_O (Sigma-Aldrich), 500 μM phosphonate-azide (prepared as described in van Bergen et al., 2023 [12]), and 25 mM sodium ascorbate (Sigma-Aldrich) in a final volume of 2 ml. Samples were incubated for 2 h at room temperature while rotating. Methanol–chloroform precipitation was performed to remove the CuAAC components, and the air-dried pellets were resuspended in 500 μl 8 M urea and sonicated in a bioruptor (Diagenode) with high amplitude for 10 min with cycles of 30 seconds on and 30 seconds off.

### Sample Processing for Digestion

Clicked and dissolved protein samples were diluted to 4 M urea with 50 mM ammonium bicarbonate (pH 8, AmBic, Sigma-Aldrich). The proteins were reduced with 4 mM DTT (Sigma-Aldrich) for 60 min at room temperature and alkylated in the dark using 8 mM iodoacetamide (Sigma-Aldrich) for 30 min. Residual iodoacetamide was quenched by adding DTT to a final concentration of 4 mM. Next, protease incubation (Pepsin Porcine, 1:50, enzyme to protein ratio, Sigma-Aldrich) was performed for 4 h at 37 °C in 40 mM HCl in a total volume of 2 ml (pH 2). Digested material was immediately desalted using 3 cc C18 Seppak cartridges (Waters) and air dried using a vacuum centrifuge.

### Dephosphorylation

Samples were dephosphorylated prior to immobilized metal affinity chromatography (IMAC) enrichment. Desalted peptides were dissolved in 1 ml 1 × CutSmart buffer (pH 8, New England BioLabs) and incubated with 50 units of alkaline phosphatase (calf intestinal, QuickCIP, New England BioLabs) overnight at 37 °C while shaking. After dephosphorylation, all peptides were again desalted using 3 cc C18 Seppak cartridges and air-dried using a vacuum centrifuge.

### Automated Fe^3+^-IMAC Enrichment

Probe-phosphonate–labeled peptides were enriched using Fe(III)-NTA 5 μl (Agilent Technologies) in an automated fashion by the AssayMAP Bravo Platform (Agilent Technologies). Fe(III)-NTA (nitrilotriacetic acid) cartridges were primed at a flow rate of 100 μl/min with 250 μl of priming buffer [0.1% TFA, 99.9% acetonitrile (ACN)] and equilibrated at a flow rate of 50 μl/min with 250 μl of loading buffer (0.1% TFA, 80% ACN). The flow through was collected into a separate plate. Dried peptides were dissolved in 200 μl of loading buffer and loaded at a flow rate of 2 μl/min onto the cartridge. Columns were washed with 250 μl of loading buffer at a flow rate of 20 μl/min, and the phosphonate-labeled peptides were eluted with 35 μl of ammonia (10%) at a flow rate of 5 μl/min directly into 35 μl of formic acid (10%). Flowthroughs and elutions were air-dried afterward, and stored at −20 °C.

### LC-MS/MS

Prior to analysis, dried peptides were dissolved in 20 μl of 2% formic acid supplemented with 20 mM citric acid (Sigma-Aldrich). Subsequently, 4 % and 45 % of the IMAC-enriched peptides were injected for the MS method optimization and dual-probe analysis, respectively. Peptides were separated by an Ultimate 3000 nanoUHPLC system (Thermo Fisher Scientific) equipped with an Aurora series column (75 μm × 25 cm, 1.6 μm, C18; Ion Opticks) heated to 50 °C by an external column oven (Sonation). The peptides were separated in the 72 min linear gradient (13.1 min 3% B, 85.1 min 30% B) at a flow rate of 400 nl/min using 0.1% FA in Milli Q as solvent A and 0.1% FA in acetonitrile as solvent B. The LC system was coupled to a trapped ion mobility quadrupole time-of-flight mass spectrometer timsTOF HT (Bruker Daltonics) via a nanoelectrospray ion source CaptiveSpray (Bruker Daltonics).

Data acquisition on the timsTOF HT was performed using TIMSControl 4.0.5.0 and Compass HyStar 6.0.30.0 (Bruker Daltonics) starting from the DDA-PASEF method optimized for standard proteomics. This method utilized a capillary voltage of 1600 V, nebulizer dry gas flow rate of 3.0 l/min at 180 °C, MS/MS target intensity of 20,000 counts, and dynamic exclusion of precursor release after 0.4 min. Singly charged peptides were excluded by an active inclusion/exclusion polygon filter applied within the ion mobility over *m/z* heatmap. Data were acquired in a range of 100-1700 *m/z* with 10 PASEF ramps (100 ms accumulation/ramp) with a total cycle time of 1.17 s. For timsTOF method optimization, the selected parameters were tested. Namely, the TIMS range of 0.6-1.6 and 0.7-1.3 Vs/cm^2^, precursor Intensity threshold of 1500 and 2500, combinations of linearly interpolated ion mobility-dependent collision energies (20, 30 eV at 0.6 Vs/cm^2^, and/or 60, 80 eV at 1.6 Vs/cm^2^), and precursor charge restriction 2+ to 5+ or 3+ to 5+ (methods A-G are summarized in Figure 1A). For subsequent concentration-dependent dual-probe experiments, method D was used (TIMS range 0.7-1.3 Vs/cm^2^, precursor intensity threshold 1500, CE 30-60 eV, and charge states 2+ to 5+).

### Database Search and Analysis

LC-MS/MS run files were searched against the human (20,375 entries) SwissProt database (version September 2020) using Fragpipe v19.1 with MSFragger 3.7, IonQuant 1.8.10, and Philosopher 4.8.1 search engine using the default settings [51]. The integrated Fragpipe contaminant database was used for filtering out contaminants. The cleavage site was set to nonspecific and a peptide length between 5 and 30 was allowed. Oxidation of methionine, acetylation of the protein N-terminus, and carbamidomethylation of cysteines were set as variable modifications. PF-06672131-phosphonate (689.20422 Da) and PF-6422899-phosphonate (632.14632 Da) adducts were also set as a variable modification on cysteine. All modifications were used in the first search. Precursor and fragment mass tolerance were set to 20 and 50 ppm respectively. The false discovery rate for PSMs and proteins was set to 1% using a target-decoy approach.

### Data analysis, Statistical Analysis, and Visualization

In the optimization for the timsTOF HT and the analysis of the characteristics of both probes, the “psm.tsv” tables were used for analysis, and all psm.tsv data was combined in a table with R. The data were graphed in GraphPad Prism 9.5.1. The quantitative dual-probe analysis was performed in Skyline-daily 22.2.1.542. The “.pep.xml” and “.d” files were loaded in Skyline-daily to enable MS1 quantification and MS/MS visualization. An ion mobility library was generated based on the results in the 10 μM probe concentration in A431 and A549 cells and corresponding peaks were integrated at similar retention times in the lower probe concentrations. Precursor peaks were filtered based on the isotope dot product (idotp) score of 0.9 in Skyline-daily. The filtered results were exported as a result table and further processed in RStudio 2023.6.2.561 [52]. Peptides without detection by a PSM in two out of three replicates at the 10 μM probe concentration were filtered out. Then, precursor peaks were filtered for presence in 2 out of the 3 replicates per condition. Afterward, the data were filtered for continuity over the concentrations on the modified peptide level (*i.e.* If a modified peptide was not found in a concentration, all values in the concentration below were filtered out). Then, to calculate the intensity per binding site, the summed MS1 peak areas of all peptides derived from the ABP binding sites were taken. Further processing and analysis of data was done in RStudio and Excel 2016 and visualization of graphs was done in Graphpad Prism 9. All spectra were exported from Skyline-daily and 3D protein modeling was conducted using UCSF ChimeraX 1.6.1 [53]. The figures were compiled and visualized using Adobe Illustrator 27.0.

## Acknowledgments

We acknowledge support from the Netherlands Organization for Scientific Research (NWO) funding the Netherlands Proteomics Centre through the X-omics Road Map program (project 184.034.019). NWO is also acknowledged for support through the VENI grant VI.Veni.202.020 (MB) and the TA project 741.018.201 (AJRH and JF).

## Conflict of interest statement

The authors declare no competing financial interest.

## Data availability

This article contains supplementary figures and supplemental data. The mass spectrometry proteomics data have been deposited to the ProteomeXchange Consortium via the PRIDE [54] partner repository with the dataset identifier PXD045864.

## Nomenclature

ABP: Activity-based probe
ABPP: Activity-based protein profiling
CID: Collision-induced dissociation
CTSC: Cathepsin C
CuAAC: Copper(I)-catalyzed azide-alkyne cycloaddition
DMAM: Dimethylaminomethyl
DPBS: Dulbecco’s phosphate-buffered saline
DUS2: tRNA-dihydrouridine synthase
EGFR: Epidermal growth factor receptor
GMPS: Guanosine monophosphate synthase
idotp: Isotope dot product
IMAC: Immobilized metal affinity chromatography
LC-MS: Liquid chromatography – Mass spectrometry
NTA: Nitrilotriacetic acid
PF131: PF-06672131
PF899: PF-6422899
PSM: Peptide spectrum match
SLC25A4/5/6: Solute carrier 25A4/5/6
SOAT1: Sterol O-acyltransferase 1
TIMS: Trapped ion mobility spectrometry
VDAC2: Voltage-dependent anion channel 2

## Notes

### Competing Interest Statement

The authors have declared no competing interest.

