## Supplemental information for "Dual-Probe Activity-Based Protein Profiling Reveals Site-Specific Differences in Protein Binding of EGFR-Directed Drugs"

**Supporting Figures and Data**

**Supplementary Figure 1.** TimsTOF analysis enables efficient and specific detection of activity-based PF131 binding sites.

**Supplementary Figure 2.** Comparative dose-dependent profiling of multiple ABPs in a single complex proteome reveals probe-specific characteristics.

**Supplementary Figure 3.** Dose-dependent site-specific target landscape reveals probe-specific target engagement in intact A549 cells.

**Supplementary Figure 4.** Concentration-dependent quantitative analysis of ABP binding to EGFR;C797 and off-target sites.

**Supplementary Figure 5.** Dose-dependent profiling of ABP binding sites.

**Supplementary Figure 6.** Quantitative analysis of ABP binding sites on VDAC1 and VDAC3.

**Supplemental Data 1.** Identifications in optimization LC-MS runs on timsTOF HT. (XLSX)

**Supplemental Data 2.** Identifications and quantifications of PF131- and PF899-bound peptides in the dual-probe PhosID-ABPP analysis. (XLSX)

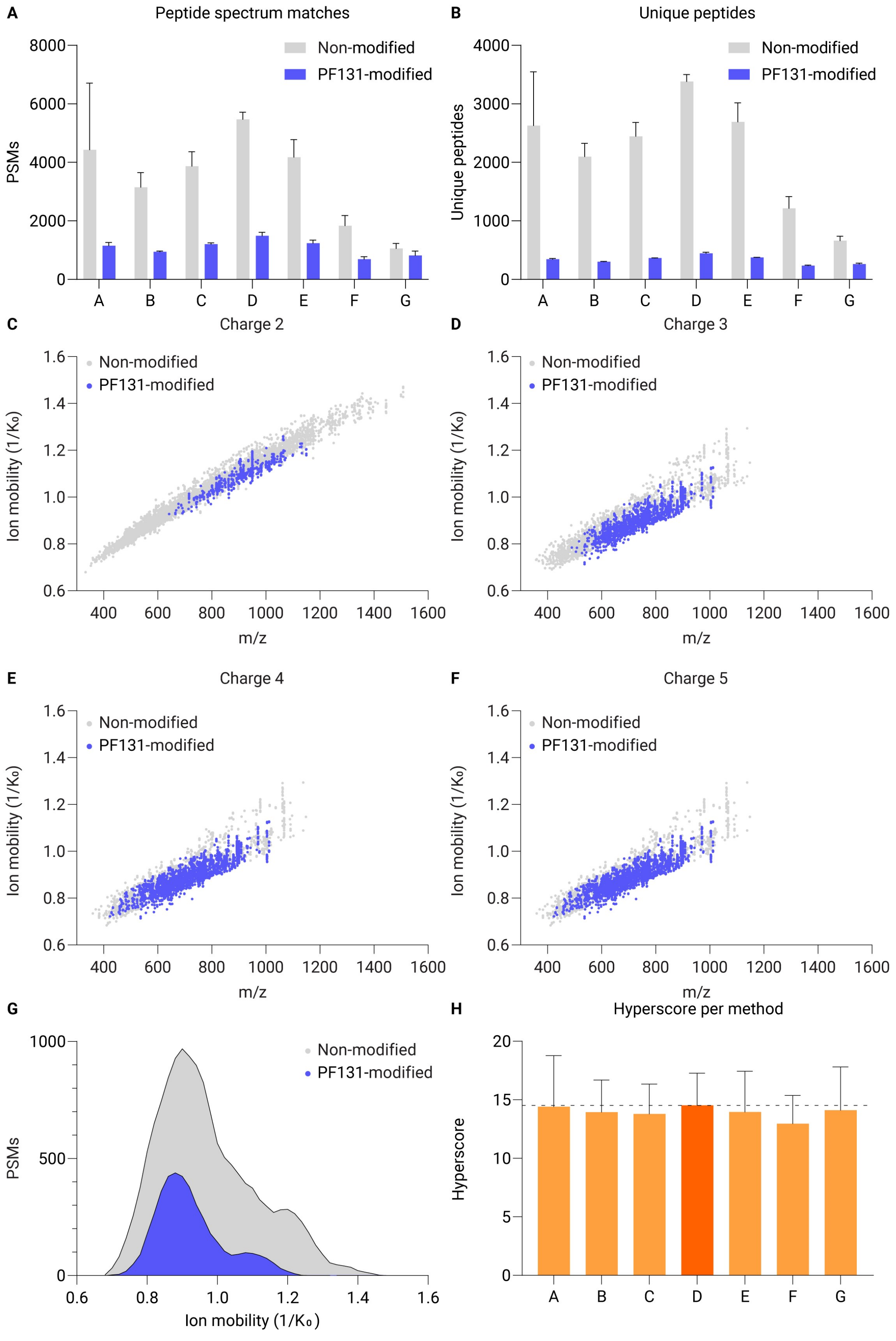

A
Peptide spectrum matches
Non-modified
PF131-modified
PSMs
8000
6000
4000
2000
0
A B C D E F G
B
Unique peptides
Non-modified
PF131-modified
Unique peptides
4000
3000
2000
1000
0
A B C D E F G
C
Charge 2
D
Charge 3
E
Charge 4
F
Charge 5
Non-modified
PF131-modified
Ion mobility (1/K₀)
1.6
1.4
1.2
1.0
0.8
0.6
400 600 800 1000 1200 1400 1600
m/z
G
Non-modified
PF131-modified
PSMs
1000
500
0
0.6 0.8 1.0 1.2 1.4 1.6
Ion mobility (1/K₀)
H
Hyperscore per method
Hyperscore
20
15
10
5
0
A B C D E F G

**Supplementary Figure 1**: TimsTOF analysis enables efficient and specific detection of PF131 binding sites. **A)** The peptide-spectrum matches (PSMs) of non-modified and PF131-modified peptides using the different methods described in Figure 1A. **B)** The number of unique non-modified and PF131-modified peptides using the different methods described in Figure 1A **C-F)** Plots displaying the ion mobility versus m/z of non-modified peptides and PF131-modified peptides detected through method A for different charge states (charge 2+ to 5+). **G)** The PSM distribution of the ion mobility for non-modified (gray) and PF131-modified (blue). **H)** Bar graph displaying the average of the hyperscores for PF131-modified peptides using the different methods described in Figure 1A. Method D shows the highest average hyperscore with a score of 14.5.

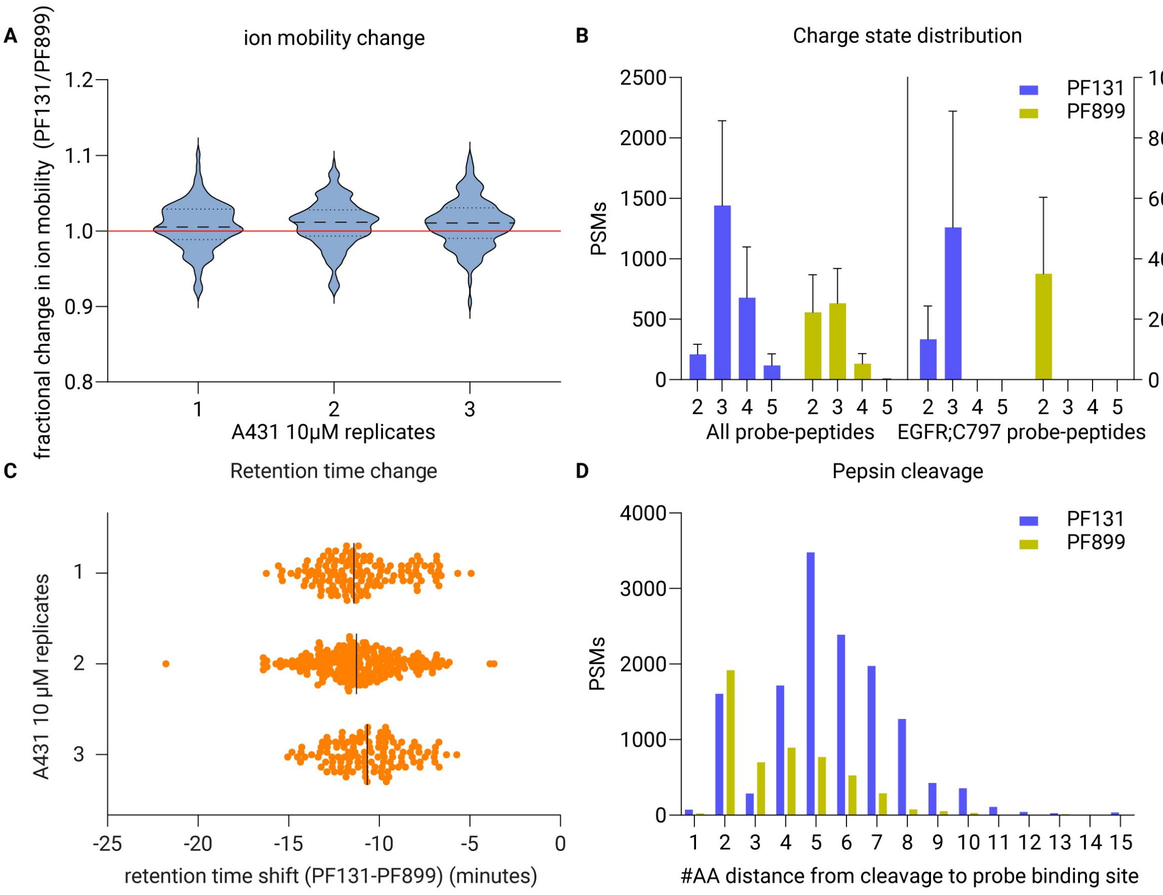


**Supplementary Figure 2:** Comparative dose-dependent profiling of multiple ABPs in a single complex proteome reveals probe-specific characteristics. **A)** Fractional change in ion mobility PF131- and PF899-labeled peptides with the same sequence in the 10 µM ABP concentration in A431 cells. The red line indicates no change in ion mobility. **B)** Peptide-spectrum matches (PSMs) corresponding to different charge states of peptides labeled with PF131 (blue) and PF899 (yellow). The left section displays the charge distribution over the entire population of ABP-labeled peptides, while the right section shows the peptides spanning the EGFR;C797 site. **C)** Shift in retention time between PF131- and PF899-labeled peptides of the same sequence in the 10 µM probe concentration in A431 cells. **D)** Distribution of peptide-spectrum matches (PSMs) corresponding to amino acid distance from the pepsin cleavage site to the ABP-bound cysteine for peptides bound to PF131 (blue) and PF899 (yellow).

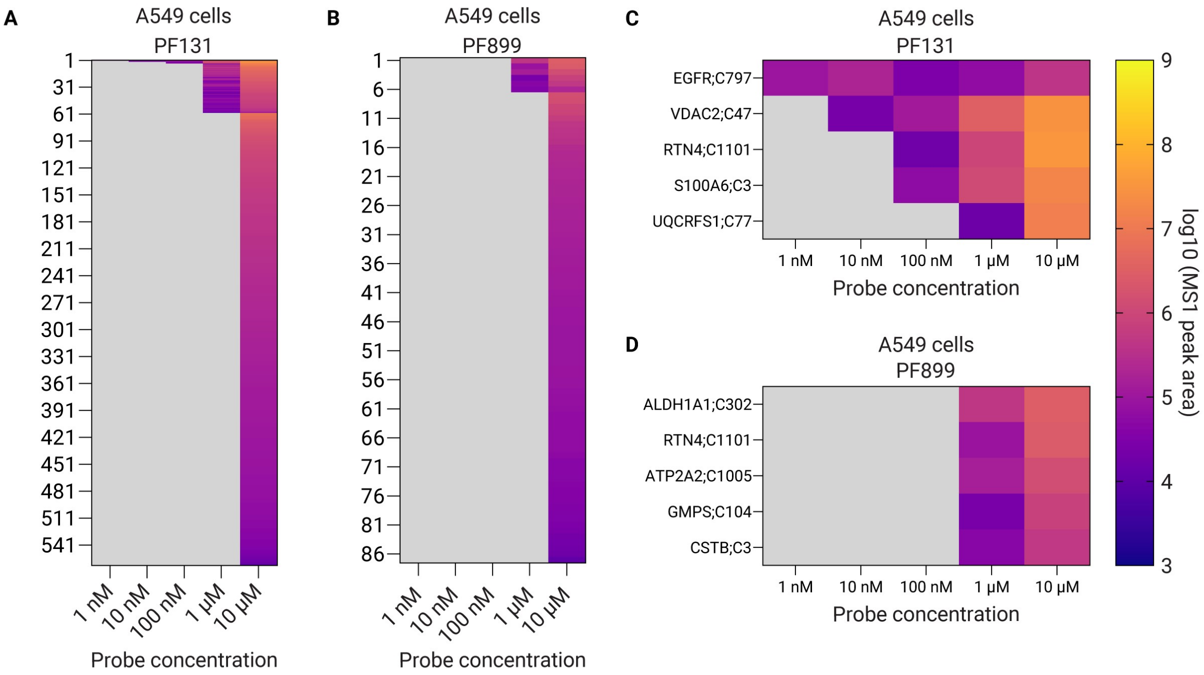


**Supplementary Figure 3:** Dose-dependent site-specific target landscape reveals probe-specific target engagement in intact A549 cells. **A)** Heatmap displaying the average aggregated MS1 peak areas per PF131 binding site at different concentrations in A549 cells. **B)** Heatmap displaying the average aggregated MS1 peak areas per PF899 binding site at different concentrations in A549 cells. **C)** A zoom of the top-5 PF131 binding sites from A. **D)** A zoom of the top-5 PF899 binding sites from B.

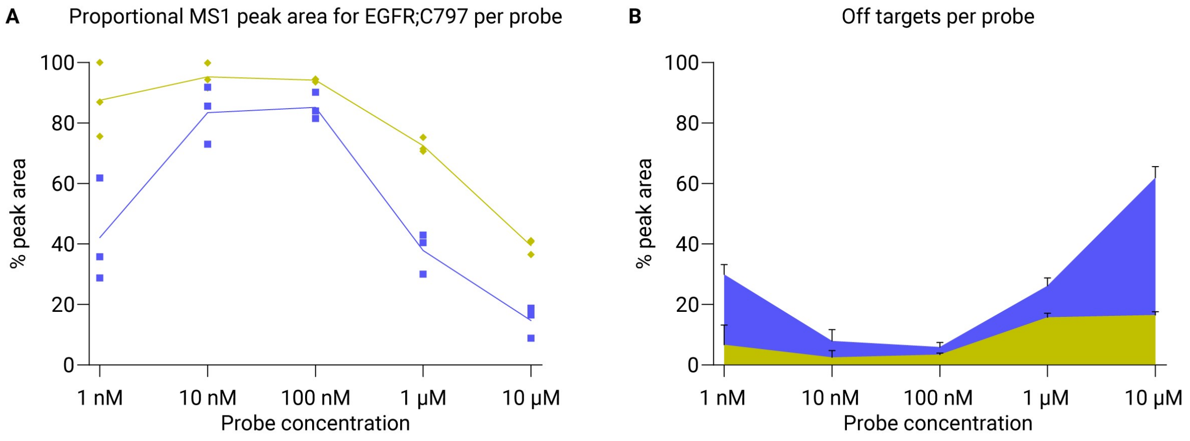


**Supplementary Figure 4:** Concentration-dependent quantitative analysis of ABP binding to EGFR;C797 and off-target sites in A431 cells. **A)** Line plot of the proportional aggregated MS1 peak areas per probe (i.e., the intensity of the specific site as a percentage of the specific probe-labeled intensity in the LC-MS experiment) for PF131 (blue) and PF899 (yellow) binding to EGFR;C797 across the concentration range. **B)** Density plot of the sum of aggregated MS1 peak areas for PF131 (blue) and PF899 (yellow) binding to any site other than EGFR;C797 across the concentration range, corresponding to the gray density in Figure 3D.

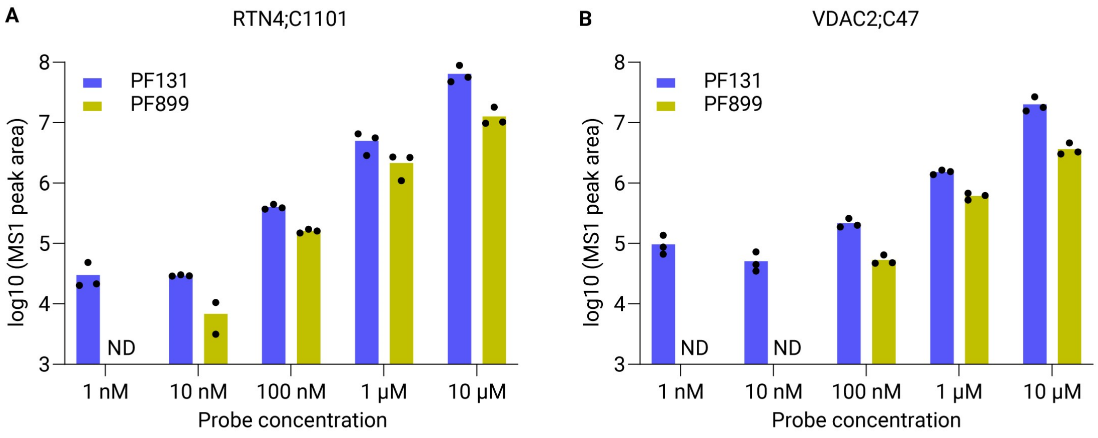


**Supplementary Figure 5**: Dose-dependent profiling of ABP binding sites in A431 cells. **A/B)** Bar graph of the aggregated MS1 peak areas for PF131 (blue) and PF899 (yellow) binding to RTN4;C1101, and VDAC2;C47 across the concentration range. ND; not detected.

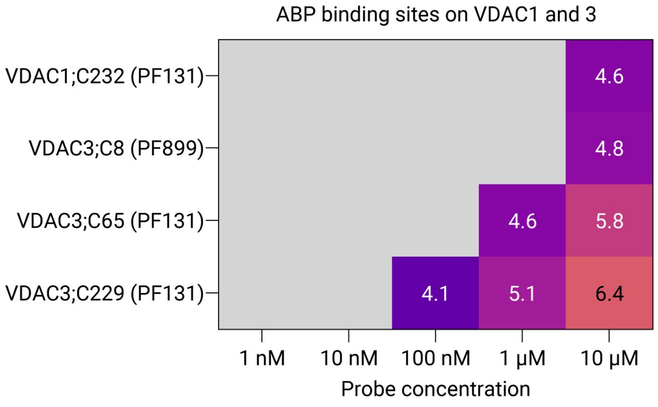


**Supplementary Figure 6:** Quantitative analysis of ABP binding sites on VDAC1 and VDAC3 in A431 cells. Heatmap displaying the log10 average aggregated MS1 peak areas per ABP binding site on VDAC1 and 3 at different concentrations in A431 cells.
